# Noise propagation in metabolic pathways: the role of growth-mediated feedback

**DOI:** 10.1101/2020.03.21.001495

**Authors:** A. Borri, P. Palumbo, A. Singh

**Affiliations:** Istituto di Analisi dei Sistemi e Informatica “A. Ruberti”, Italian National Research Council (IASI-CNR), Via dei Taurini, Roma, Italy. Email address; Department of Biotechnologies and Biosciences, University of Milano-Bicocca, Milan, Italy 20126. E-Mail address; Department of Electrical and Computer Engineering, Biomedical Engineering, Mathematical Sciences, Center for Bioinformatics and Computational Biology, University of Delaware, Newark, DE USA 19716. E-Mail address

**Keywords:** Metabolic pathways, Enzymatic Reactions, Systems Biology, Feedback

## Abstract

Metabolic networks are known to deal with the chemical reactions responsible to fuel cellular activities with energy and carbon source and, as a matter of fact, to set the growth rate of the cell. To this end, feedback and regulatory networks play a crucial role to handle adaptation to external perturbations and internal noise. In this work, a cellular resource is assumed to be activated at the end of a metabolic pathway, by means of a cascade of transformations. Such a cascade is triggered by the catalytic action of an enzyme that promotes the first transformation. The final product is responsible for the cellular growth rate modulation. This mechanism acts in feedback at the enzymatic level, since the enzyme (as well as all species) is subject to dilution, with the dilution rate set by growth. Enzymatic production is modeled by the occurrence of noisy bursts: a Stochastic Hybrid System is exploited to model the network and to investigate how such noise propagates on growth fluctuations. A major biological finding is that, differently from other models of metabolic pathways disregarding growth-mediated feedback, fluctuations in enzyme levels do not produce only local effects, but propagate up to the final product (hence to the growth rate). Furthermore, the delay provided by the cascade length helps in reducing the impact of enzymatic noise on to growth fluctuations. Analytical results are supported by Monte Carlo simulations.

## I. Introduction

Fluctuations in growth rate are known to be responsible for phenotypic heterogeneity, although the mechanisms behind them are still matter of investigation [19], [20]. A large effort has been spent to investigate such phenotypic heterogeneity since it is supposed to be involved in cellular growth control and cancer initiation (see [12] and references therein). Within this framework, metabolism has recently gained interest since the intrinsic noise in gene expression and enzymes accumulation has been shown to propagate to growth rate fluctuations through metabolic fluxes, according to singlecell experiments [9], [13], [22].

In this note a metabolic pathway is considered (see the scheme in Fig. 1), dealing with a cellular resource *X*_0_ required to undergo a set of *p* functional modifications before to exert cellular growth control as *X_p_*. Such modifications are triggered by an enzyme *E*. The activated resource *X_p_* indirectly controls, in turn, the dilution of all molecular players, by means of the growth rate which is linearly related to *X_p_* accumulation.

**Fig. 1.**
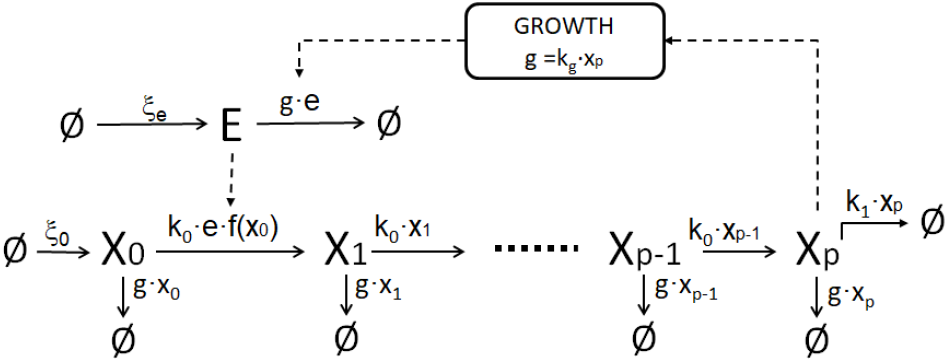
Schematic figure of the metabolic pathway under investigation. Continuous lines refer to reactions; dashed lines refer to control actions.

Stochastic fluctuation in metabolic pathways has been matter of investigation in a wide range of literature, aiming at understanding the implications of gene expression stochasticity in cellular regulation and phenotypic diversity [15], [10], [4]. In [11] the analytical approach of a mathematical model exploiting Chemical Master Equations to deal with the cascade of chemical reactions showed that, for a linear metabolic pathway, fluctuations in different nodes of the pathway are uncorrelated and, consequently, fluctuations in enzyme levels have just local effects and do not propagate through the line of metabolic transformations.

The metabolic network here investigated differs from the framework adopted in [11], since we account for the feedback exerted by the final product that controls, by means of growth modulation, the dilution rate of all involved species. A similar interplay between enzyme, metabolites and growth rate has been investigated in [3], where the cellular metabolic resource *X* was not subject to a functional modification. Moreover, in [3] growth influenced only the enzyme dilution, whilst here all molecular players share the same dilution rate inherited from growth, according to a more biologically meaningful assumption.

A Stochastic Hybrid System (SHS) is exploited in order to model the network of biochemical reactions [8]. All molecular players are treated as copy numbers. Enzymatic production is supposed to occur in a discrete stochastic fashion: the enzyme copy numbers are increased by a discrete amount each time a burst occurs. Within any two consecutive bursts of enzymatic production, the other molecular players are supposed to vary continuously according to an ordinary Differential Equation (oDE) system.

The goal of the manuscript is to investigate how noise propagates from enzyme to growth, by means of the metabolic transformations. To this end, the Coefficient of Variation is exploited in order to quantify the noise affecting molecular players fluctuations [11], [14], [2]; the analysis of the cross-correlation coefficient is also exploited to understand whether noise fluctuations propagate from the enzyme to cellular growth, or vice versa. In both cases, first- and second-order moments are required. Because of the ODE nonlinearities, finite-order moments equations are not provided in closed form [8], thus preventing any analytical computations except those involving moment closure techniques (e.g. the ones developed in [17]). To overcome such a problem, here we resort to linearizing the ODE nonlinearities around the steady-state solution that is proven to be the unique stationary equilibrium point.

Results show that the longer is the length of the cascade, the less the noise in enzyme production impacts on the final product fluctuations, hence proving that the delay provided by the cascade length helps in limiting growth fluctuations. Moreover, the cross-correlation analysis highlights the occurrence of an apparent delay from enzyme to growth fluctuations (and not vice versa, similarly to [3]), with such delay increasing with the cascade length as well. Further results achieved at a fixed cascade length show that fluctuations at enzymatic level propagate through the pathway, differently from cases without the feedback exerted by growth, like the one developed in [11].

The note is organized as follows. Section II introduces the model setting and Section III addresses the computation of first-order moments. Section IV is devoted to the computation of second-order moments and autocorrelation functions. Section V illustrates stochastic simulations aiming at validating the theoretical results. Section VI offers concluding remarks.

## II. Model setting

Consider the metabolic pathway depicted in Fig. 1. Copy numbers of resources and enzyme will be accounted for, denoting them with x_0_, x_1_,…, x_1_ and e. The enzymatic production is the only noise source: it is assumed to occur in bursts, with the burst size *μ* a random variable taking values in {1,2,…}. Then, whenever a burst occurs, e is updated as follows

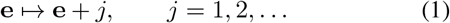

with a propensity 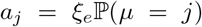. According to past literature [18], [2], [7], such probability is chosen to comply with a geometric distribution

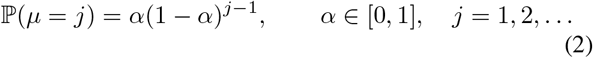

providing an average burst size 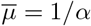.

Between any two bursts, the state variables evolve according to an ODE describing all species dynamics by properly setting the reaction rates of all chemical reactions.

The reaction rate of the first transformation *X*_0_ ↦ *X*_1_:

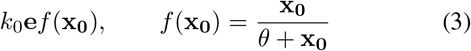

is controlled by the enzyme in a linear fashion, whilst *X*_0_ enters according to a saturating function. This kind of saturating functions are often exploited in molecular control mechanisms, such as model transcripts production rates controlled by transcription factor accumulation [1]. All other transformations in the cascade (*X_i_* ↦ *X*_*i*+1_, *i* = 1,…, *p*−1) have a reaction rate linearly dependent of only x_i_:

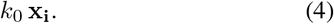

All species are subject to dilution, with the dilution rate provided by the growth rate *g* which in turn is proportional to *X_p_* copy number:

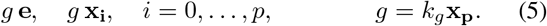

Indeed, *X_p_* is supposed to modulate the growth rate by properly influencing the cell metabolism. The growth rate closes the loop on the enzyme dynamics, since it rules the dilution of *E*. Actually, the growth rate rules the dilution of all other molecular players. Besides, *X_p_* is also cleared out according to a linear clearance rate

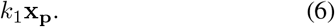

In summary, the ODE is written as follows

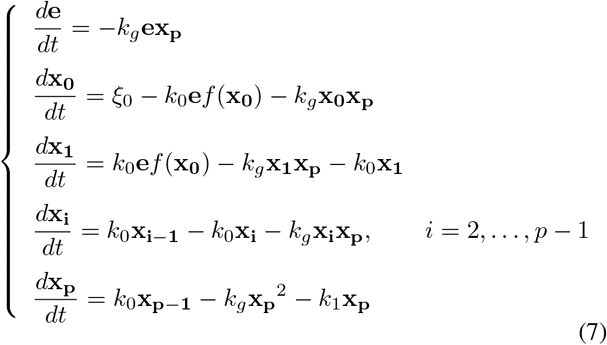

and it will be shortly denoted by

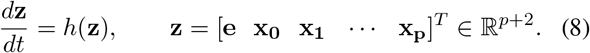

## III. First-order moment equations

The expected value dynamics of any scalar transformation 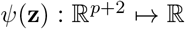 of the state of the SHS described by the continuous ODE dynamics in (7)–(8) and endowed with the discrete resets (1) obeys to the following equation [8]:

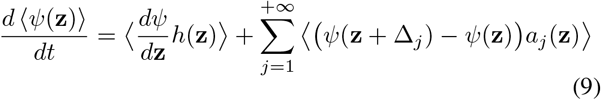

where Δ_*j*_, *j* = 1,2,… is the displacement on z because of reaction *j* in (1). This formula is exploited to compute the dynamics of any order moment. For instance, with respect to the first-order moments, we have:

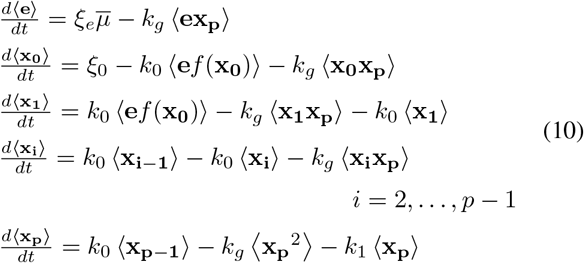

Because of the nonlinear terms, first-order moment equations cannot be written in a closed form [8], therefore we resort to the linear approximations of the oDE nonlinearities around the stationary point

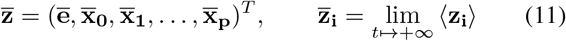

provided by:

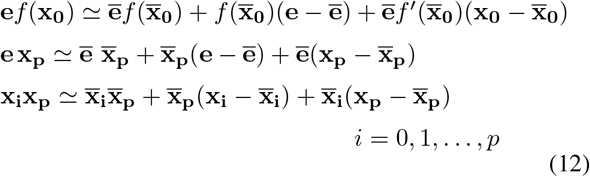

Thus, first-order moment equations are written, after computations, according to the following linear system

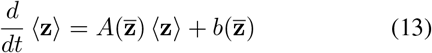

Matrix 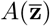 and vector 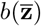 are explicitly reported in Appendix.

By properly exploiting the fact that linearizations are achieved around the stationary point, this equilibrium has to comply with the following constraints

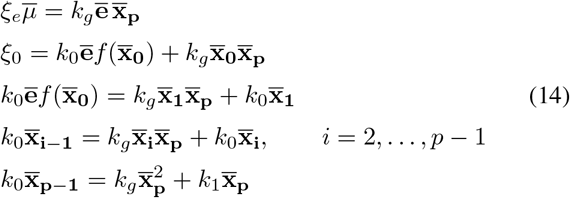

### Theorem 1

There exists a unique, positive real solution for the nonlinear algebraic system (14), for any choice of positive model parameters.

*Proof:* As a preliminary step it will be proven that, for

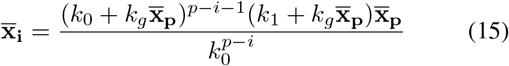

The proof comes from mathematical induction. Indeed, (15) is true for *i* = *p* − 1, since, by straightforward substitution, we readily obtain the last constraint in (14). Now, let (15) be true for a given *i*. Then from the penultimate constraint in (14), one gets:

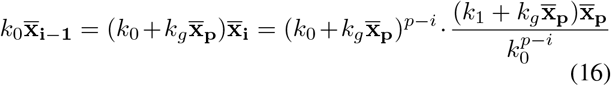

from which it straightforwardly comes that (14) is true also for *i* − 1.

By properly manipulating the second and third equations in (14) one has:

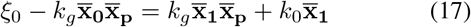

from which, after substituting in (17) eq. (15) computed for *i* = 1, we have also 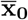 as a function of 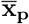:

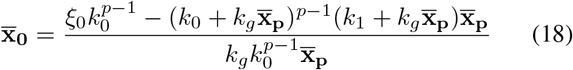

This relationship will be referred to in the sequel as 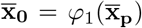.

On the other hand, also 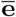 can be easily written as a function of 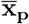 from the first equation of (14):

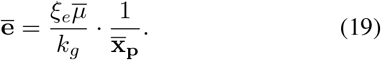

Therefore, after substituting (19) and eq. (15) computed for *i* = 1 in the third constraint of (14), by suitably exploiting the saturating function (3), one has the following further relationship between 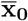 and 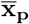:

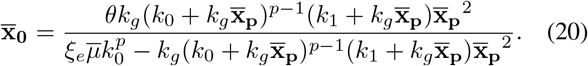

This relationship will be referred to in the sequel as 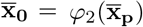.

In summary, the biologically meaningful stationary solutions are provided by the positive real intersections of the curves (18)-(20) in the 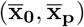 plane. Regards to 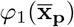, it is monotonically decreasing in the positive orthant, with

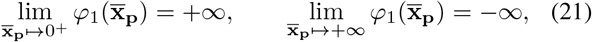

so that it has a unique positive value for which it vanishes. On the other hand, the second curve is monotonically increasing in the positive orthant, with

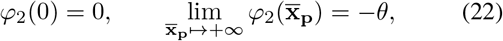

with a vertical asymptote in the unique point that vanishes the denominator of *φ*_2_(·). As a matter of fact, there exists a unique intersection of the two curves in the positive orthant.

## IV. Second-order moment equations

According to what anticipated in the Introduction, we exploit the following definition of metabolic noise (e.g. for enzyme or growth fluctuations) provided by the squared Coefficient of Variation [11], [14], [2]:

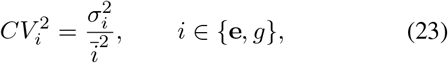

where *σ_e_, σ_g_* are the standard deviations of e and *g*.

Second-order moments are computed according to (9) [8]:

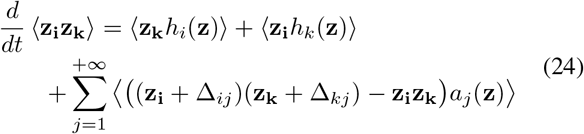

where Δ_*ij*_, *i* = 1,…,p + 2, *j* = 1, 2,… is the displacement on z_i_ because of reaction *j*.

Because of the linearizations (12), the stationary values of the second order moments equations provided by (24) are the solutions of a linear system, whose formal expression is achieved according to straighforward, though cumbersome computations, and is not here reported. The order of the system, by properly exploiting the simmetries, is:

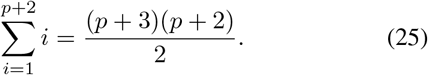

In order to evaluate how noise fluctuations in the enzyme impact in growth, we compute the stationary crosscorrelation

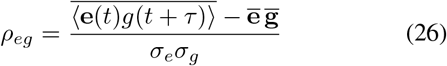

where 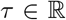 refers to the noise propagation delay and the overbars denote the stationary average values so that, for example:

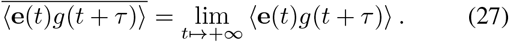

Because of the linear relationship between *g* and x_p_, (5), one has:

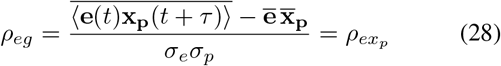

with *σ_p_* being the standard deviation of x_p_. Correlation analysis is carried out according to the same approach developed in [16], [3] so that, for *τ* ≥ 0 we take advantage of

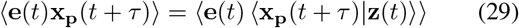

Then, by exploiting the explicit solution of the linear equation in (13), we have, provided that *A*^−1^ exists,

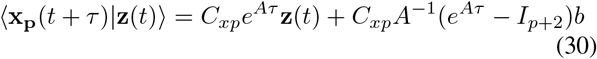

where *C_xp_* = [*O*_1×(p+1)_ 1], *I*_*p*+2_ is the identity matrix in 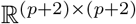. By substituting (30) in (29), we have

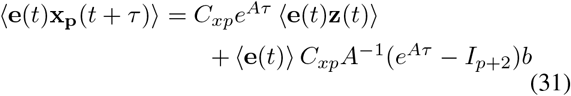

and, at steady-state

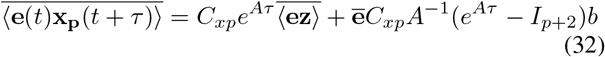

where 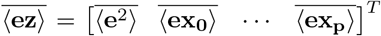 are part of the stationary second-order moments dealt with at the beginning of the Section.

In case of negative *τ*:

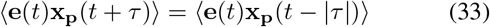

so that, at steady-state, it is

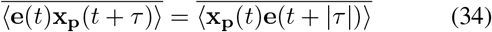

**TABLE I.**
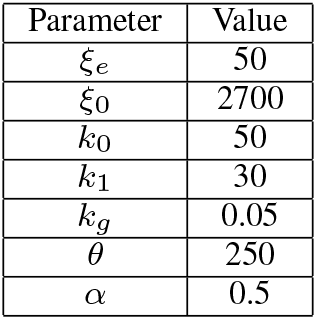
Model parameters

The computation of 〈x_p_(*t*)e(*t* + |*τ*|)) follows the same steps developed before:

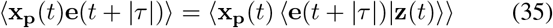

with

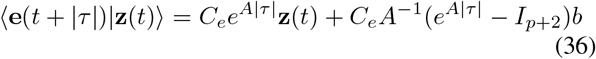

where *C_e_* = [1 *O*_1×(p+1)_] so that

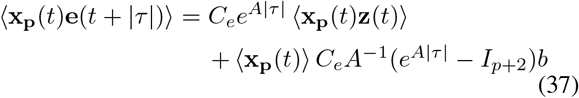

and, at steady-state

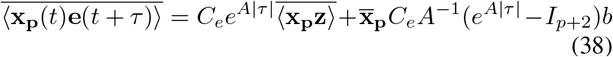

where 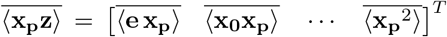 are part of the stationary second-order moments dealt with at the beginning of the Section.

## V. Simulation results and discussion

Analytical solutions concerning both first- and second-order moments achieved according to linear approximations of the nonlinear terms of the ODE. Therefore, results here reported are validated by Monte Carlo stochastic simulations for the SHS. These simulations are performed by means of the tau-leaping algorithm [6], by also exploiting the ergodic properties of the underlying stochastic process [21]. In this setting, the step selection has been chosen equal to 0.001 seconds within an overall simulation time of 2000 seconds. The model parameters are reported in Table I.

A first set of simulations aims at showing how the cascade length impacts on noise propagation. To this end, all model parameters have been fixed as the ones reported in Table I. Both analytical results and Monte Carlo simulations show that by increasing *p*, the steady-state average values of enzyme and final product copy numbers have opposite effects: stationary enzyme values increase, whilst stationary final product values decrease, see Fig. 2. The wiring of the network helps in understanding this behavior: by increasing the length of the cascade, nontrivial amount of substrate is stuck on intermediate values, thus reducing the final product accumulation; on the other hand, a reduced final product accumulation weakens the dilution of all molecular players, including the enzyme, thus allowing a greater enzyme accumulation. Fig. 3 reports how the noise associated to growth fluctuations varies for varying values of *p*, and we find out that such *CV*s decrease with *p*. That means, the delay provided by the cascade length helps in limiting growth fluctuations.

**Fig. 2.**
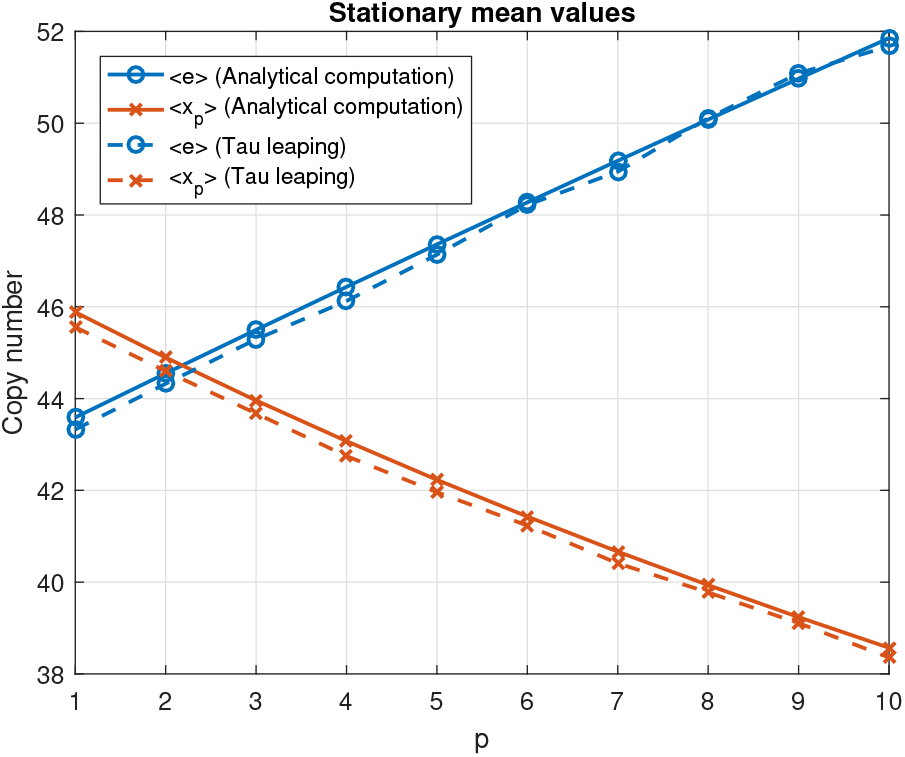
Stationary mean values for variable cascade length (up to *p* = 10)

**Fig. 3.**
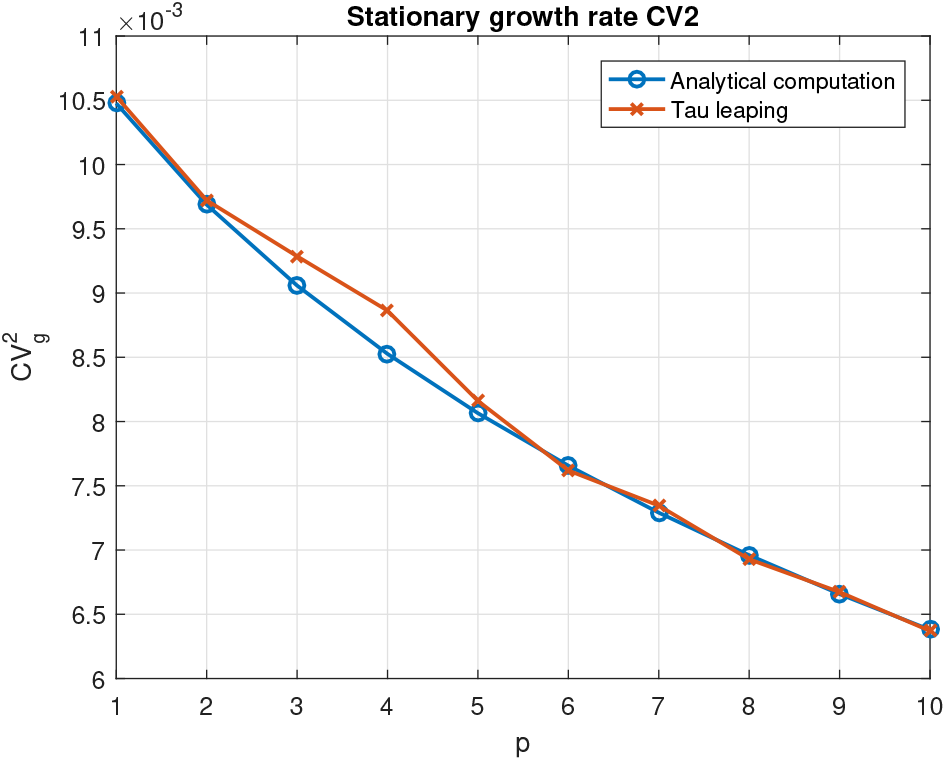
Stationary values for the *CV*^2^ of the growth rate, for variable cascade length (up to p = 10)

When plotting the cross-correlation *ρ_eg_*(*τ*) (see Fig. 4), we find out that there exists an apparent delay showing that current enzyme expression correlates better with growth at some time later. This fact is coherent with experimental results reported in [22], thus explaining that growth fluctuations occur because of the noise in the enzyme expression, and not vice versa. This results had also been achieved in a similar analysis, although obtained according to a simplified network of players. Indeed, the present model allows to investigate also how the length of the cascade impacts on such delay. Fig. 4 shows that the delay increases with *p*.

**Fig. 4.**
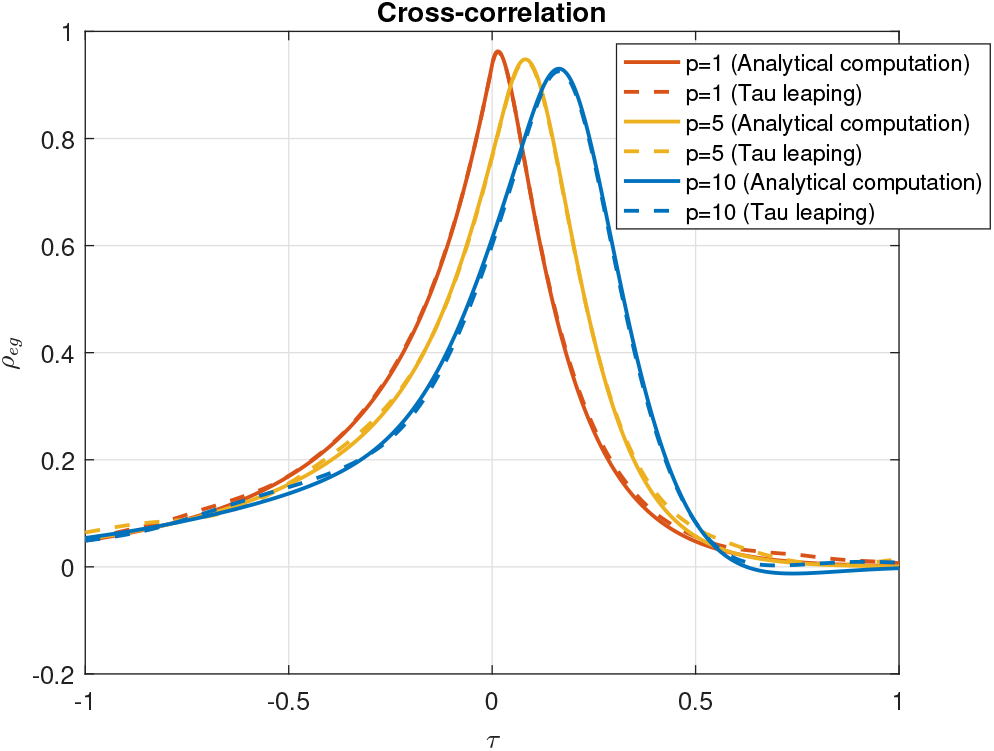
Cross-correlation *ρ_eg_* for different cascade lengths

A second set of simulations has been carried out for a fixed value of the cascade length (*p* = 10), by properly varying the enzymatic noisy production. Indeed, by varying the parameter *α* of the enzyme burst geometric distribution in (2) (with respect to the nominal value in Table I), it is possible to tune the variance of the incoming noise for the enzyme. It is worth to notice that, by varying *α* ∈ [0,1], one obtains a wide range of burst size fluctuations and, as a matter of fact, different levels of enzymatic noise.

In order to make a fair comparison, we properly tuned the model parameters to keep fixed stationary average values in spite of different burst sizes. Looking at the constraints in (14), it comes out that *α* affects only the first equation (by means of 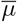), therefore we have properly modified *ξ_e_* to have the product 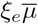 fixed for any value of *α*. More in details:

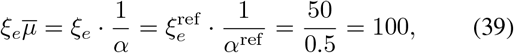

so that *ξ_e_* = 100*α*, with reference values for 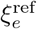 and *α*^ref^ taken from Table I. Fig. 5 reports how growth fluctuations levels vary according to enzyme noise levels (in both cases quantified by their *CV*^2^). It is apparent that, by increasing the stochastic enzymatic fluctuations, also the growth noise increase. This fact enhances the role of the growth-mediated feedback exerted by the final product accumulation since, without the feedback, it has been shown in [11] that fluctuations in enzyme levels do not propagate through the pathway.

**Fig. 5.**
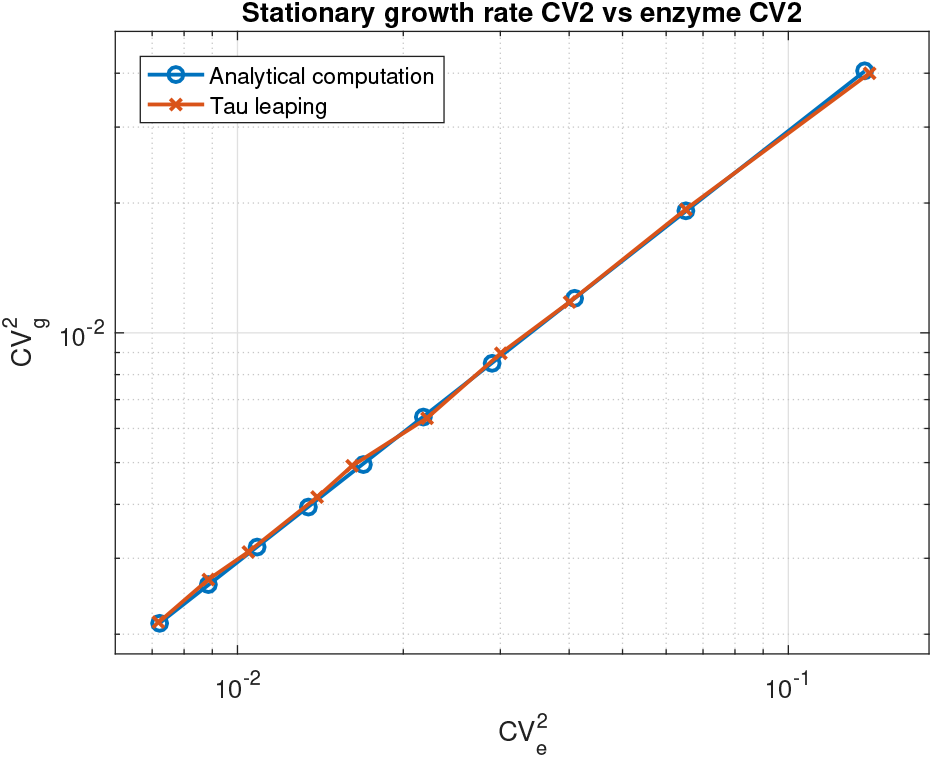
Stationary growth rate 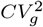 vs enzyme 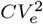

## VI. Conclusions

In this note, we have investigated a metabolic pathway where a cellular resource undergoes a cascade of functional modifications, triggered by an enzyme, before exerting cellular growth control. Approximate moment computations and correlation functions aim at investigating how noise propagates from enzyme production through growth; in a biological context, understanding the fluctuations in growth rate is of topic importance, since noise is known to be responsible for phenotypic heterogeneity, which is supposed to be involved, for example, in cancer initiation and growth. The analytical computations are based on a Stochastic Hybrid System framework, and further validated via approximate stochastic simulations. Previous works [11] have shown that stochastic fluctuations do not propagate from enzymatic activity in the intermediate states of a linear metabolic pathway. Instead, here we show that, by accounting for a growth-mediated feedback from the final product accumulation, enzymatic noise levels have no more only local effects, with the length of the cascade playing a crucial role in noise propagation.

## Appendix

With respect to eq.(13), the nontrivial entries of matrix 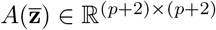 are

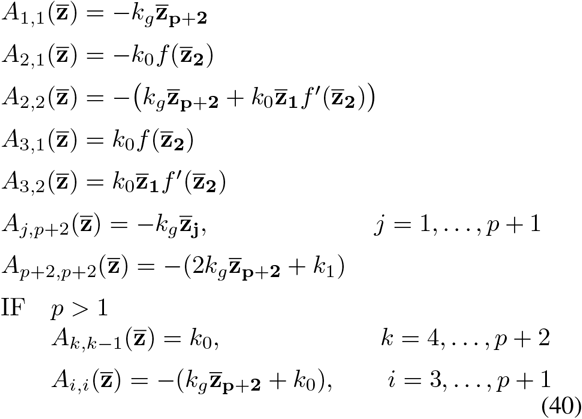

whilst, regards to 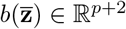, we have:

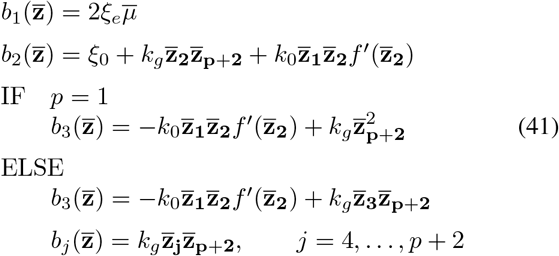

